# Deciphering the Genetic Architecture of Sorghum Grain Oil Content via Lipidome-Integrated Genome-Wide Association Analysis

**DOI:** 10.64898/2026.03.12.711187

**Authors:** Deepti Nigam, Sarah Metwally, Fang Chen, Yinping Jiao

**Affiliations:** Institute of Genomics for Crop Abiotic Stress Tolerance (IGCAST), Department of Plant and Soil Science, Texas Tech University, Lubbock, Texas, 79409 USA; Center for Biotechnology and Genomics, Texas Tech University, Lubbock, Texas, 79409 USA

**Keywords:** Sorghum grain oil, Lipidomics, Lipidome-wide association study (lGWAS), Triacylglycerol (TAG), Stacked alleles, High-oil genotypes, Crop biofortification

## Abstract

Grain oil content and composition are complex quantitative traits that shape cereal grain quality and nutritional value. Sorghum (*Sorghum bicolor*), a heat- and drought-adapted C₄ crop essential for global food and feed security, remains insufficiently characterized with respect to grain lipidome diversity and its genetic architecture. Here, we integrated population-scale whole-grain lipidomics with genome-wide association studies (GWAS) in 266 sorghum accessions. Lipidome profiling revealed extensive natural variation in triacylglycerols (TAGs), accompanied by coordinated shifts in phosphatidylcholines (PCs) and phosphatidylethanolamines (PEs), explaining 87% of population-level differences in total grain oil. Lipidome-wide GWAS identified approximately 1.6 million significant variant-trait associations and resolved 55 loci linked to plastidial fatty acid synthesis, TAG assembly, lipid transport, and membrane remodeling. These loci, many undetected in previous GWAS of bulk oil content, demonstrated the increased mapping resolution achieved through lipidomics. Integration with metabolic gene clusters revealed significant enrichment of lipid-associated variants within terpene and saccharide-terpene biosynthetic clusters, indicating coordinated genetic regulation between central lipid metabolism and specialized metabolic pathways. Variants within these clusters explained more than 50% of the variance in measured grain oil content and exhibited additive effects of favorable alleles. Haplotype analyses further identified 27 elite sorghum accessions and 12 linked markers for marker-assisted improvement of sorghum grain oil. These findings elucidate the multilayered genetic architecture of sorghum grain lipid diversity and showcase the value of large-scale lipidomics integrated with GWAS for accelerating C₄ crop grain quality improvement.

## INTRODUCTION

Grass crop grains are not only the foundation of global food security but also a rich source of diverse fatty acids that play vital roles in human health and industrial applications (GUO *et al*. 2022; VERMA *et al*. 2023). Oils derived from grains consist primarily of saturated (SFAs), monounsaturated (MUFAs), and polyunsaturated fatty acids (PUFAs), each influencing physiology in distinct ways (DAMUDE AND KINNEY 2008). Beyond providing high-quality energy, these oils supply essential nutrients and bioactive compounds with important health benefits (VUKIC AND VUJASINOVIC 2023; SHARMA *et al*. 2024). For example, PUFAs are integral components of membrane phospholipids in specific tissues and serve as precursors for signaling molecules such as prostaglandins (WIKTOROWSKA-OWCZAREK *et al*. 2015; DAS 2018; HARWOOD 2023). In recent years, fatty acids with specific pharmacological or functional properties have gained considerable attention from both consumers and industries, highlighting their growing economic and health relevance (KAUR *et al*. 2014; CALDER 2015). Improving the nutritional quality of crop grain oils stands as a major goal for crop breeding, which requires a deeper understanding of the biological and genetic factors that shape fatty acid composition (TONG *et al*. 2025).

Recent advances in lipidomics and genomics now make it possible to link biochemical diversity to genetic variation, providing new insight into how lipid biosynthesis is regulated in cereal grains (BATES 2016; LI *et al*. 2019; CAO *et al*. 2023; TIOZON JR *et al*. 2025). Fatty acid synthesis primarily occurs in plastids, whereas subsequent lipid assembly and modification are carried out in the endoplasmic reticulum. Key enzymes, including acetyl-CoA carboxylase (ACCase), fatty acid desaturases (FADs), and diacylglycerol acyltransferases (DGATs), play pivotal roles in determining both the total oil content and the composition of TAGs and membrane lipids (BATES 2016; YE *et al*. 2020; XIAO *et al*. 2022; ZHOU *et al*. 2025). Insights from cereal crops, particularly maize (*Zea mays*), where genome-wide association studies (GWAS) have dissected the complex genetic control of kernel oil biosynthesis (LI *et al*. 2013; LI *et al*. 2019; FANG *et al*. 2021). For example, the high-oil maize accessions have been used to genetically map loci affecting carbon partitioning, fatty acid elongation, and TAG assembly, including *DGAT1-2*, *OLE1*, and various transcription factors regulating oil biosynthesis (LI *et al*. 2010; DE ABREU E LIMA *et al*. 2018; LUO *et al*. 2023).

Among cereal grains, sorghum (*Sorghum bicolor*) stands out for its exceptional resilience to drought and heat, making it a strategic crop for sustainable agriculture due to its high drought tolerance and low input requirements (TARDIEU *et al*. 2018; FONTANET-MANZANEQUE *et al*. 2025). As a staple food for an estimated 500 million people worldwide, sorghum plays a critical role in global food security (KHALIFA AND ELTAHIR 2023; REZAEI *et al*. 2023). Although sorghum is starch-rich, its lipid fraction contributes significantly to grain quality, energy density, and functional properties relevant to food and industry (ABAH *et al*. 2020; TANWAR *et al*. 2023). Enhancing the oil content and composition of sorghum grain offers significant opportunities to improve nutritional value, diversify industrial applications, and bolster farmer livelihoods(AFIFY *et al*. 2012). Nevertheless, the understanding of sorghum grain lipid metabolism remains limited, particularly with respect to the genetic factors controlling lipid diversity and accumulation. Most previous studies have focused primarily on total oil content, leaving minor but functionally important lipid classes, such as sphingolipids and sterols, largely unexplored (ZHU *et al*. ; HEUPEL *et al*. 1986; MEHMOOD *et al*. 2008). Recent advances in lipidomics have started to fill this knowledge gap, as demonstrated by a nontargeted lipidomics analysis that identified previously unreported fatty acid esters of hydroxy fatty acids in sorghum grain and uncovered pronounced cultivar-specific differences in lipid composition (NATH *et al*. 2024). Genetic engineering of sorghum using maize lipid-regulatory genes has demonstrated that coordinated up-regulation of lipid biosynthesis can substantially increase oil accumulation, underscoring opportunities for translational discovery and allele stacking to enhance grain oil content (VANHERCKE *et al*. 2019). These studies have focused primarily on total oil concentration, with relatively few efforts aimed at characterizing the diversity of individual lipid species or identifying the genetic determinants underlying this variation *(MEHMOOD et al. 2008; MOREAU et al. 2016; KIMANI et al. 2020; COSSIO et al. 2025).* The lack of high-resolution, comprehensive lipidomic, wide population genetic study has hindered progress in identifying genetic resources for improving sorghum grain oil content and quality.

To address this knowledge gap, we performed UHPLC-MS-based lipidomic analysis of whole grains from 266 sorghum association panel (SAP) accessions, quantifying the broad spectrum of grain lipid species. Integrating these data with genome-wide association studies (GWAS) enabled us to identify approximately one million variants controlling lipid biosynthesis and accumulation pathways. This integrative approach provides new insight into the molecular mechanisms shaping sorghum grain composition, paving the way for targeted breeding of nutritionally enhanced and industrially valuable sorghum varieties.

## RESULTS

### Lipidome composition of sorghum grain

To capture lipidome variation of sorghum grain in the diversity population, we conducted high-throughput untargeted lipidomics on 266 SAP accessions. In total, 1,060 lipid species were detected, including 626 positive-mode electrospray ionization (ESI+) features and 434 negative-mode electrospray ionization (ESI-) features; 528 of these species were detected in >75% of the accessions in the population and annotated into six major super-classes (Supplementary Fig. 1A; Supplementary file 1).

Fatty acyls were the most abundant super-class (152 species; 28.8%), followed by glycerophospholipids (GPLs; 142 species; 26.9%), glycerolipids (GLs; 117 species; 22.2%), sterol lipids (71 species; 13.4%), prenol lipids (42 species; 8.0%), and sphingolipids (4 species; 0.8%). GPLs were dominated by phosphatidylethanolamines (PE) and phosphatidylcholines (PC), whereas GLs were led by triacylglycerols (TAGs) with minor inputs from galactolipids and diacylglycerols. Correlation analysis of compound-level abundance profiles showed that overall lipid composition clearly separated sorghum accessions into distinct groups (Supplementary Fig. 1B, Supplementary file 2).

At the individual lipid species level, TAGs were the most abundant class, followed by PCs and PEs, whereas phosphatidylinositols (PI), phosphatidylglycerols (PG), and galactolipids showed lower levels (Fig. 1A). Ranking the top 20 lipid species confirmed this dominance, with phosphatidylcholine 14:0-20:3 being most abundant, followed by lysophosphatidylcholine 18:1 and multiple triacylglycerols containing common fatty acids 16:0, 18:1, and 18:2 (Fig. 1B). A complementary PCA including genotype scores and lipid loadings yielded a similar structure, with the first two components explaining the majority of variance (Supplementary Fig. 2A) and a subset of lipid species exhibiting long loading vectors that drive separation among genotypes, indicating that a relatively small number of abundant and highly variable lipids underpins the major axes of lipidomic differentiation (Supplementary Fig. 2B). Together, these results indicate that sorghum grain lipidome variation is driven primarily by a few dominant TAGs and membrane phospholipids (PCs/PEs), which constitute the primary structural components of seed lipid composition.

**Figure 1.**
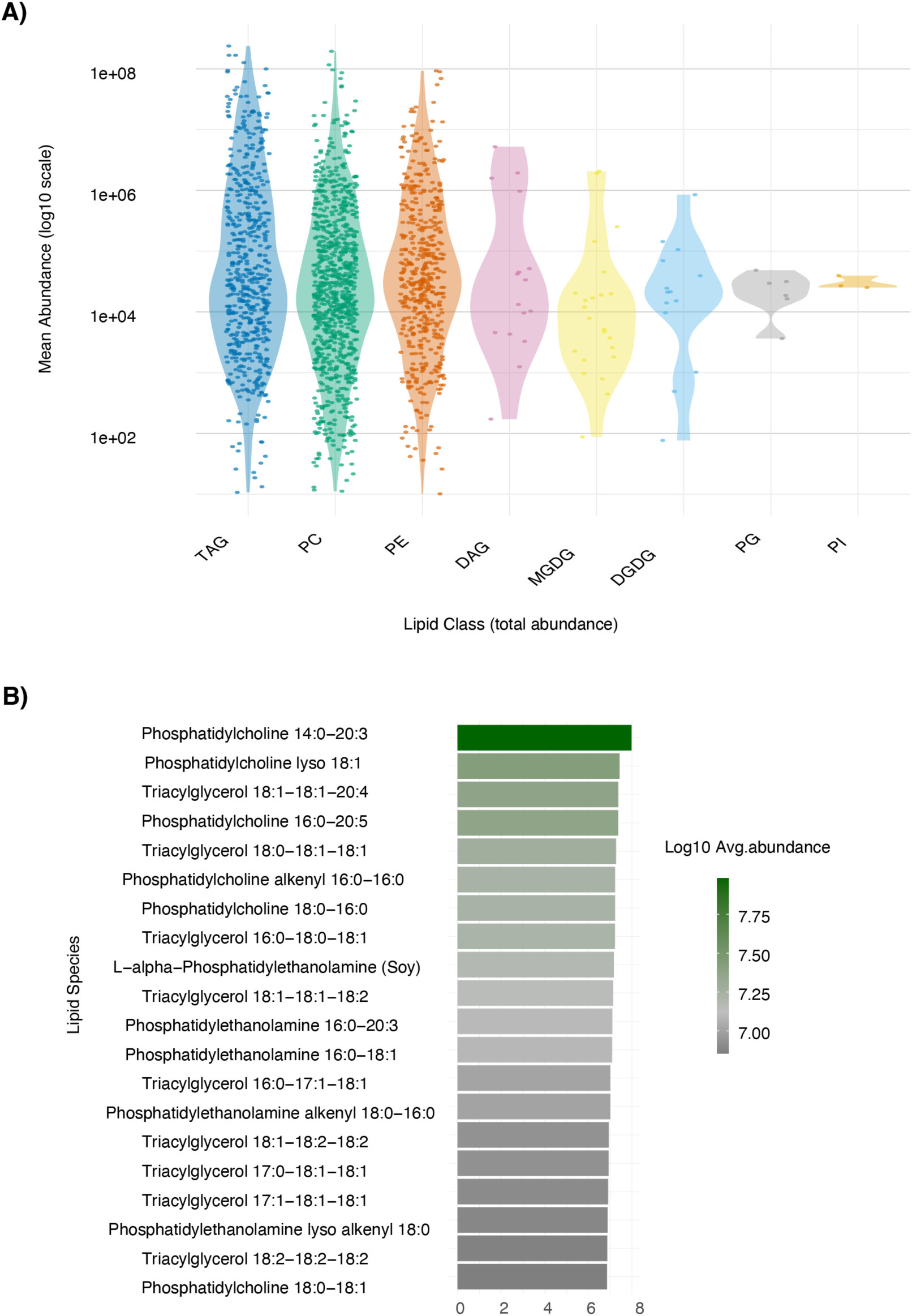
Landscape of lipid classes and species in sorghum grain. **(A)** Distribution of mean lipid abundances across major lipid classes, including phosphatidylcholine (PC), phosphatidylethanolamine (PE), phosphatidylglycerol (PG), phosphatidylinositol (PI), triacylglycerols (TAG), diacylglycerols (DAG), digalactosyldiacylglycerols (DGDG), and monogalactosyldiacylglycerols (MGDG), measured by UHPLC-MS in 266 sorghum accessions. Each point represents a lipid species, and violin plots display the distribution of mean log₁₀ abundances within each lipid class. **(B)** The 20 most abundant lipid species detected across the sorghum association panel. Bars represent the average log₁₀ abundance of individual lipid species, with color intensity corresponding to relative abundance.

### TAG-driven variation in grain oil content across the SAP

The SAP is known to harbor substantial diversity in grain oil content (RHODES *et al*. 2017). Using the 528 lipid species detected in more than 75% of accessions, we characterized lipidome diversity across the panel. These 528 species span the major lipid classes-TAG, DAG, PC, and PE-which collectively represent crude oil content. Shannon diversity analysis, which integrates the richness and evenness of lipid species within each class, showed that TAGs, PCs, and PEs together account for ∼87% of total lipidomic variation across all 266 accessions (Fig. 2A).

**Figure 2.**
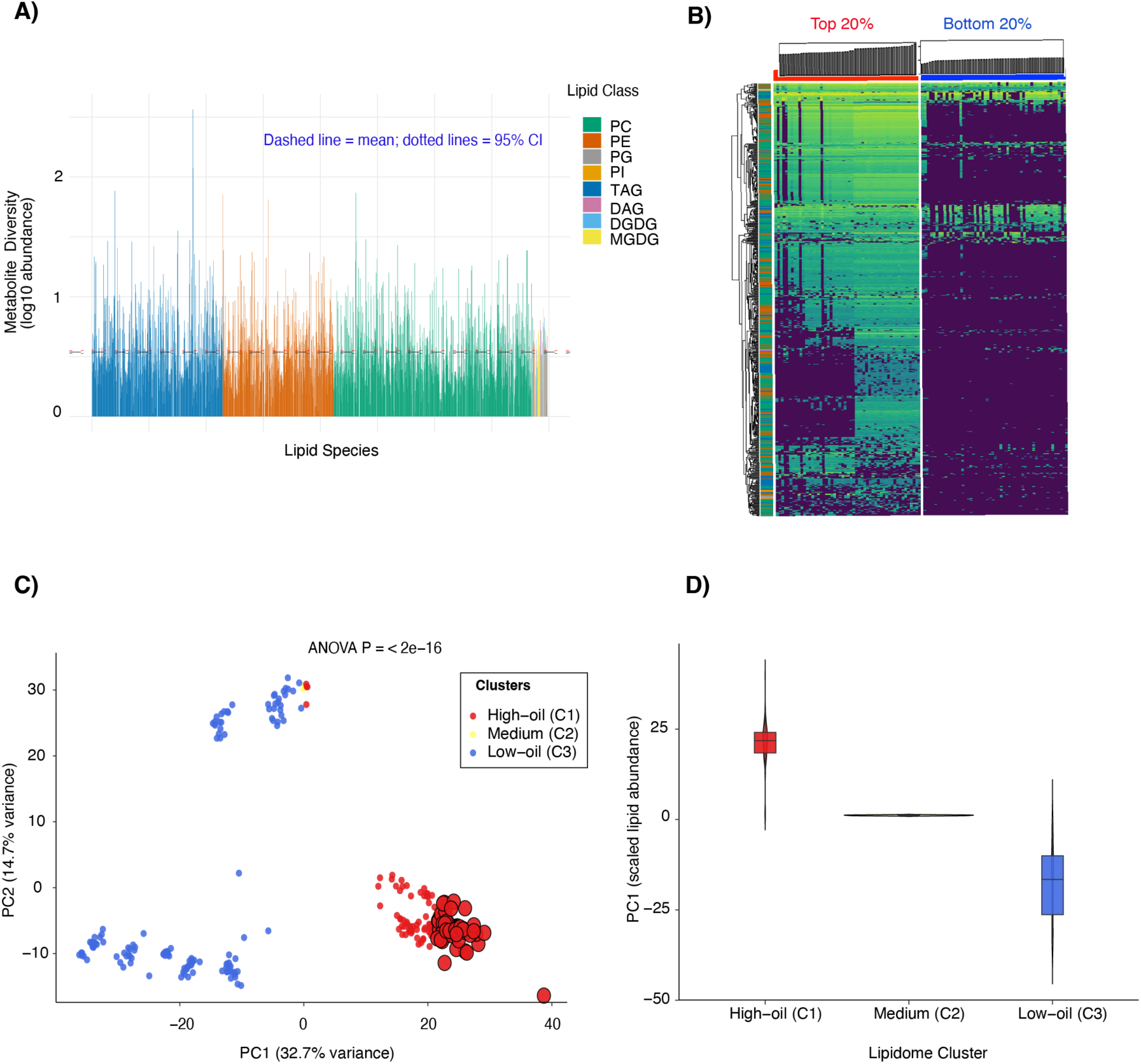
Sorghum grain lipidome variation. **(A)** Distribution of log-transformed metabolite abundances across individual lipid species grouped by lipid class (PC, PE, PG, PI, TAG, DAG, DGDG, and MGDG). The dashed horizontal line represents the overall mean abundance, and the dotted lines denote the 95% confidence interval. (B) Heatmap of hierarchical clustering based on scaled lipid abundances comparing the top 20% and bottom 20% oil-content accessions among 266 sorghum accessions. Rows represent lipid species and columns represent accessions. (C) Principal component analysis (PCA) of lipid abundances showing the distribution of sorghum accessions along PC1 and PC2. Each point represents an accession colored by lipidome cluster (high-oil [C1], medium [C2], and low-oil [C3]). The percentages indicate variance explained by each principal component. (D) Distribution of PC1 values across lipidome clusters (C1-C3).

We further classified the 266 sorghum accessions into three groups based on lipid abundances: high-oil (n = 124), medium-oil (n = 3), and low-oil (n = 139). Hierarchical clustering of log₁₀-transformed lipid abundances resolved three distinct lipidomic groups with strong separation evident in the dendrogram structure (Supplementary Fig. 3A; ANOVA across clusters, *P* < 10⁻¹⁵). Approximately 85% of detected TAG species displayed significant cluster-specific accumulation patterns (P < 0.01), whereas minor lipid classes remained uniformly low across groups. Principal component analysis further supported these patterns: an ordination based on all lipid species and another weighted by mean abundance both placed TAGs and PCs near the center of lipidomic space, whereas PE, PI, PG, DG/DAG, and MG/MGDG occupied more peripheral positions (Supplementary Fig. 3B and C).

Comparison of the top and bottom 20% of sorghum accessions ranked by total lipid abundance showed that high-lipid accessions accumulated ∼3.5-fold higher median levels of specific TAGs, PCs, and PEs (Fig. 2B). Principal component analysis further confirmed this partitioning of lipidomic variation: PC1, derived from scaled lipid abundances, explained 32.7% of the total variance and clearly separated high- and low-oil accessions, with high-oil accessions (C1) exhibiting strongly positive PC1 values and low-oil accessions (C2) displaying negative PC1 values (Fig. 2C; Supplementary file 3). Consistent with this pattern, the distributions of PC1 scores differed significantly among lipidome clusters (Fig. 2D).

Together, these results indicate that variation in grain oil content among SAP accessions is driven primarily by differences in TAG accumulation, with accompanying shifts in PC and PE levels that reflect altered glycerolipid partitioning. High-oil lipidomic profiles (C1) therefore represent promising targets for breeding, and the substantial heterogeneity observed within the high-oil fraction (Fig. 2D) highlights additional opportunities to further optimize oil biosynthesis and turnover through genetic improvement.

### GWAS of the sorghum grain lipidome

Given the strong correlations and distinct clustering patterns among lipid species (Supplementary Fig. 1B), we performed both univariate and multivariate GWAS (FERNANDES *et al*. 2021; SMITH *et al*. 2022), using 38,873,896 published SNPs with allele frequency >=0.05 (BOATWRIGHT *et al*. 2022) (Fig. 3A). Univariate GWAS was conducted for each of 528 individual lipid species, whereas multivariate GWAS was performed jointly with six lipid clusters, defined by shared headgroup class and acyl-chain composition (triacylglycerols, PCs/PEs, sphingolipids, galactolipids, phosphatidylinositols, and minor lipid classes) (Fig. 3A). After correcting for population structure with 58 principal components that explained 90% of lipidome variance, univariate analyses identified 919,846 variant-lipid associations surpassing the Bonferroni threshold (*P* < 1 × 10⁻⁸), while multivariate analyses identified 647,543 variant cluster associations at the same threshold, with genomic inflation factors ranging from 0.98 to 1.15 (Supplementary file 4).

**Figure 3.**
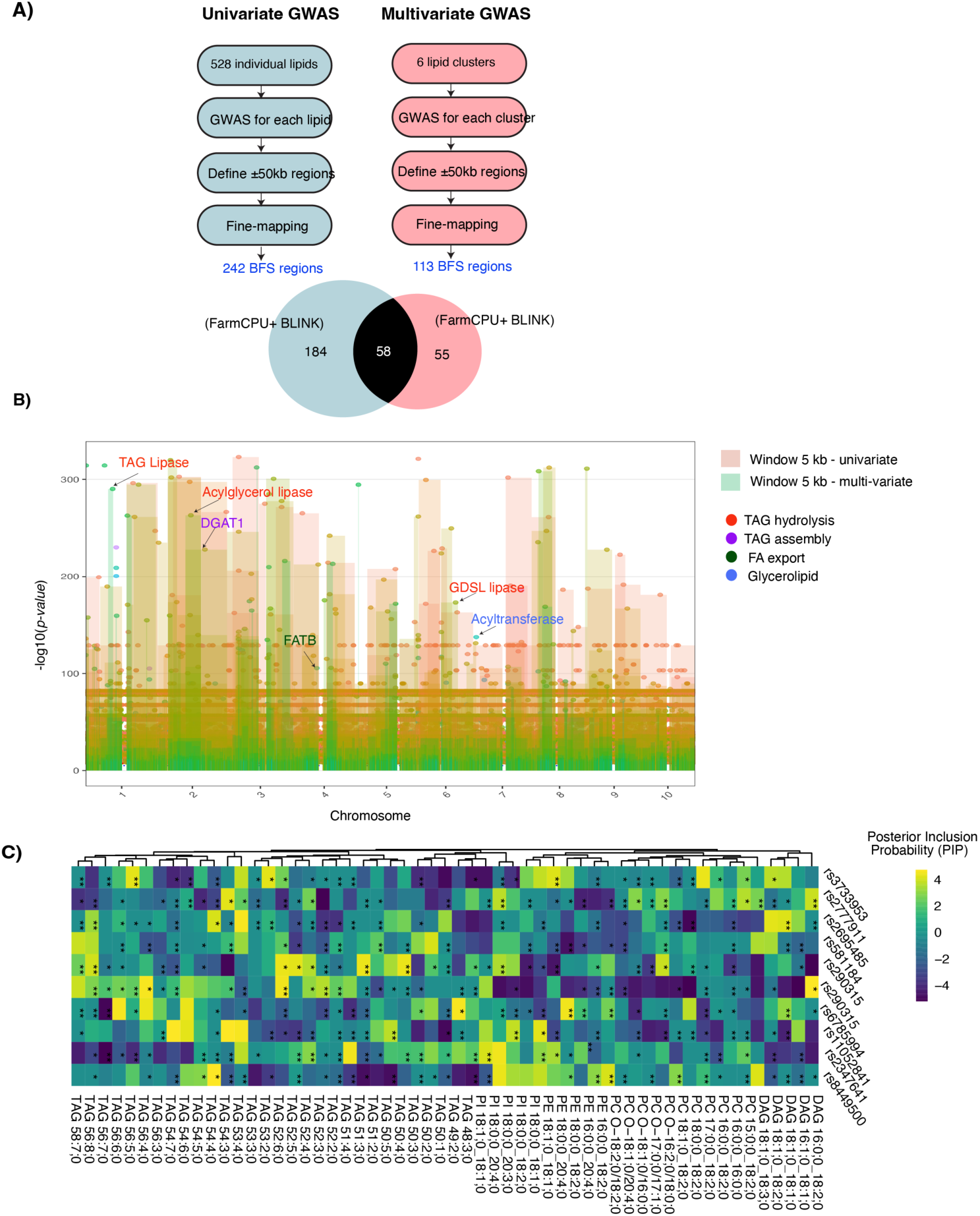
Single- and multi-trait GWAS of sorghum grain lipid variation. (A) Workflow comparing single-trait (univariate; blue) and multi-trait (multivariate; pink) GWAS using the Bayesian Multi-Trait (BNF) model (univariate: 240 regions; multivariate: 105 regions), with comparison of loci detected (Venn: 70 shared, 170 univariate-specific, 35 multivariate-specific).(B) Distribution of association signals from the two approaches across chromosomes (windows ±5 kb around candidates; univariate in light gray, multivariate in orange).(C) Posterior inclusion probabilities (PIP) from the multivariate model across lipid species (rows: SNPs; columns: lipid species).

To compare the two approaches, we defined lead variants for each trait and merged ±50 kb windows around these leads using linkage disequilibrium (r² ≥ 0.1) (Supplementary file 5). The analysis identified 355 significant association regions: 242 detected exclusively by univariate GWAS, 113 exclusively by multivariate GWAS, and 58 shared between the two approaches (Fig. 3A and B). Significant associations for TAG, PE, and PC were observed across all chromosomes (Supplementary Fig. 4A). Several genomic regions showed associations with more than one lipid class, whereas other regions appeared to be lipid-class specific. Multivariate analysis was found to yield lower local false sign rates (LFSR)-the posterior probability that an estimated effect has the wrong sign-than univariate analysis. The largest LFSR improvements occurred for variants affecting multiple lipid clusters, as shown by the downward shift of multivariate points below the diagonal (Supplementary Fig. 4B).

Posterior inclusion probabilities (PIP) from the multivariate model, ranging from 0 (no evidence of association) to 1 (strong evidence)-showed variable patterns across the six lipid clusters (OTTENSMANN *et al*. 2023) (Fig. 3C). Some loci exhibited associations specific to individual clusters, whereas others showed associations across multiple lipid classes, suggesting partially shared genetic influences on the grain lipidome. One example is a locus detected only by the multivariate analysis on chromosome 10, located within the sorghum diacylglycerol acyltransferase 1 gene region (*DGAT1*; *Sobic.010G170000*, Chr10: 50,099,835-50,118,712 bp). The lead SNP at 50,106,710 bp (G/T) was located within the gene and showed relatively high PIP across several TAG species. This region colocalized with a previously reported grain oil QTL and showed both positive and negative effects across lipid species (Fig. 3C, D), consistent with the role of DGAT1 in TAG biosynthesis biosynthesis(ZHANG *et al*. 2009). Collectively, these results demonstrate that multivariate GWAS improved the power to detect loci in the sorghum grain lipidome; therefore, all downstream analyses focus on the 55 association signals identified by the multivariate approach (Supplementary file 5).

Within this refined set, eight harbor high-confidence candidate genes (*P* < 10^−1^) in core lipid-biosynthesis pathways (Supplementary file 5). Four independent loci contain genes encoding rate-controlling enzymes of plastidial de-novo fatty acid synthesis (FAS II): *Sobic.002G151000* (enoyl-[acyl-carrier-protein] reductase I, FAB1), and a tandem array of three paralogues (*Sobic.001G231400, Sobic.001G231600, Sobic.001G231700*) encoding 3-oxoacyl-[acyl-carrier-protein] reductase (KAR/FABG). Two loci contain *Sobic.001G251200,* encoding a P4-ATPase phospholipid flippase required for membrane lipid asymmetry and lipid droplet formation, and *Sobic.002G134100*, encoding fatty acyl-CoA reductase, the committed step in cuticular wax synthesis. Notably, these eight loci were not previously detected in GWAS of total oil content (RHODES *et al*. 2017), underscoring the power of lipidome-based GWAS to uncover previously unreported regulators of cereal grain lipid content and composition.

To assess lipid pathway connections to broader metabolism, we mapped GWAS loci with previously defined metabolic gene clusters (MGCs) (Nigam et al. 2025). These 38 clusters comprise 4,347 metabolite biosynthetic genes and represent physically co-localized groups of enzymes that function together in co-regulated biosynthetic pathways (Supplementary File 6 and Supplementary Fig. 5). We identified 158 lipidome-associated variants overlapping 31 MGCs, including 7 harboring TAG/DAG GWAS variants within terpene/saccharide-terpene clusters (23.4% of total), suggesting coordinated terpenoid-lipid regulation (Supplementary Fig. 4).

Among these loci, 34 SNPs exceeded the stringent GWAS significance threshold (P ≤ 5 × 10⁻⁸) and were located within ±50 kb of terpene or saccharide-terpene gene clusters (Supplementary Fig. 5). These regions contained 26 candidate genes enriched for lipid-related metabolic functions (Supplementary Fig. 6). Enriched functional categories included glycerophospholipid and glycerolipid metabolism, ether lipid metabolism, sphingolipid metabolism, and fatty acid synthase activity, indicating the involvement of multiple lipid biosynthetic pathways (Supplementary File 6). This set was prioritized for downstream polygenic modeling because it combined strong GWAS signals with physical proximity to pathway-related genes and enrichment for lipid-associated functions. Such pathway-constrained selections improve predictive power and biological interpretability over genome-wide SNPs, as shown in other sorghum grain and oil QTL studies (KICHAEV *et al*. 2019; KUMAR *et al*. 2023; VOLPATO *et al*. 2023).

### A haplotype stacking strategy to increase total oil content in sorghum

Grain oil content in crops is a quantitatively inherited, polygenic trait governed by multiple loci with small to moderate effect (MACE *et al*. 2013; MORRIS *et al*. 2013; WALLACE *et al*. 2014). Guided by the pathway-based prioritization described above, we constructed a Polygenic Lipid Score (PLS) for each of the 266 accessions by summing favorable allele dosages across the 34 prioritized SNPs, weighted by their GWAS effect sizes (β) on total grain oil content (Fig. 4A). The overall analytical framework used to identify and integrate favorable lipid-associated alleles is summarized in Supplementary Fig. 7. Favorable alleles were defined as those associated with increased lipid abundance, corresponding to positive β coefficients in the GWAS model (BERNARDO 2008; WALLACE *et al*. 2014; BOYLES *et al*. 2017). The PLS provides a quantitative measure of the cumulative genetic contribution of lipid-associated loci to oil accumulation. The sorghum accessions were divided into five equal PLS quantiles (Q1-Q5; Fig. 5B). Accessions in the highest quantile (Q5) exhibited, on average, 24% greater oil concentration than those in Q1, with the five highest- and lowest-oil accessions occupying opposite distribution tails relative to the population mean ± SD (Fig. 4B, C). The PLS explained over half of the variation in measured oil content (R² = 0.545; Fig. 5C), indicating substantial additive genetic contribution to the trait.

**Figure 4.**
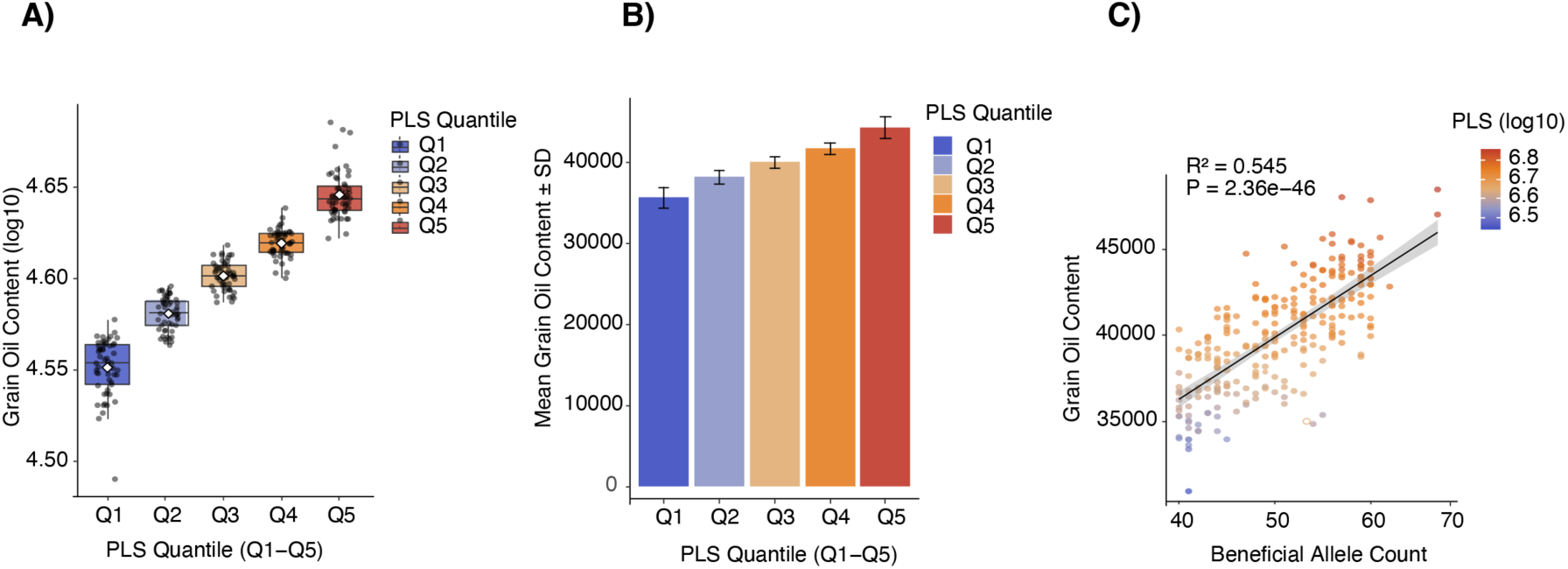
Genetic contribution of lipid-associated loci to sorghum grain oil accumulation. (A) Distribution of grain oil content across Polygenic Lipid Score (PLS) quintiles (Q1: lowest PLS; Q5: highest PLS). Boxes indicate interquartile ranges, center lines indicate medians, and points represent individual accessions (colored by quintile). (B) Mean grain oil content (± SD) for accessions grouped by PLS quintiles. (C) Beneficial alleles count vs. grain oil content across 266 sorghum accessions (points colored by PLS z-score).

**Figure 5.**
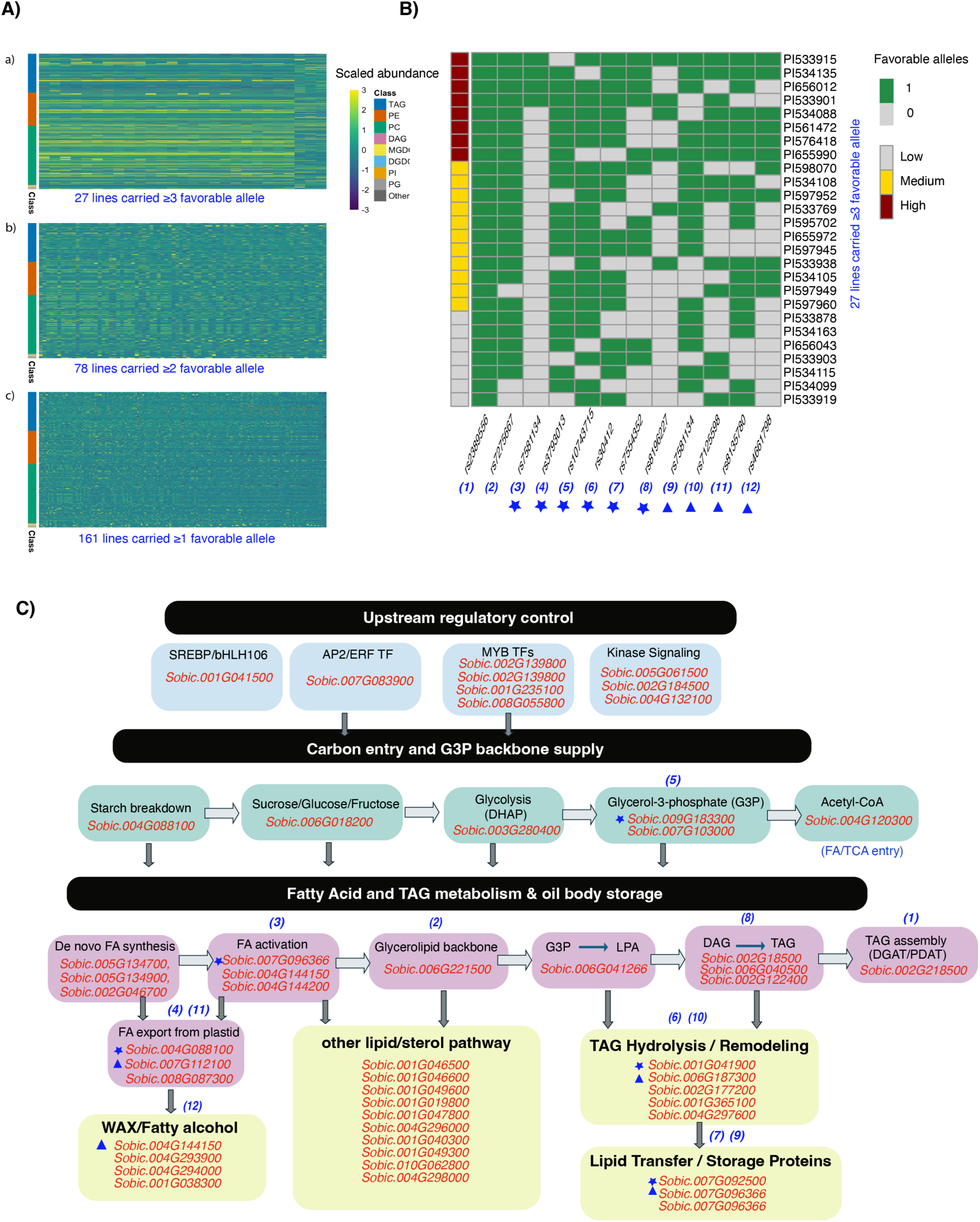
Core sorghum accessions and candidate genes associated with grain oil content. (A) Heatmaps showing scaled lipid abundances in sorghum accessions carrying different numbers of favorable alleles across the 12 selected loci. Panels display accessions carrying ≥3 favorable alleles (top), ≥2 favorable alleles (middle), and ≥1 favorable allele (bottom). Lipid species are grouped by class, and values represent scaled abundances. (B) Distribution of favorable alleles across the 12 core SNP loci in the 27 accessions carrying ≥3 favorable alleles. Green cells indicate the presence of favorable alleles (1), and gray cells indicate absence (0). Accessions are annotated by oil content category (low, medium, and high). (C) Candidate genes associated with the 12 prioritized loci mapped onto metabolic pathways related to upstream regulatory control, carbon entry and glycerol-3-phosphate backbone supply, fatty acid synthesis, glycerolipid backbone formation, TAG assembly, lipid remodeling, and lipid storage.

Based on the effects of these QTLs on total oil content, 12 core SNPs were selected from the 34 analyzed loci as candidate breeding targets for increasing grain oil content in sorghum. These 12 prioritized SNPs were chosen for their largest GWAS effect sizes (β > 0.3), central roles in the fatty acid to triacylglycerol (FA→TAG) pathway, and high minor allele frequency (> 0.1). Across these 12 loci, 161 sorghum accessions carried ≥1 favorable allele, 78 harbored ≥2, and 27 accumulated ≥3 favorable alleles (Fig. 5A; Supplementary Fig. 8; Supplementary File 8). Haplotype analysis across the 12 SNPs revealed multiple allele combinations with varying numbers of favorable alleles, and network relationships among haplotypes indicated gradual accumulation of beneficial variants (Supplementary Fig. 8A and B). Linkage disequilibrium patterns among these SNPs suggested partial independence among loci, consistent with their combined contribution to oil content variation (Supplementary Fig. 8C).

Allele stacking exhibited a positive relationship with lipid abundance: sorghum accessions with ≥3 favorable alleles showed substantially higher total oil content (mean 5.8%, range 5.2-7.1%) compared to those carrying 0-1 favorable allele (mean 3.9%; Wilcoxon rank-sum test, *P* < 2.2 × 10⁻¹⁶) (Fig. 5A). This analysis identified 27 sorghum accessions as multi-locus carriers (harboring ≥3 favorable alleles across the 12 loci) (Fig. 5B). Importantly, these 27 accessions fully overlapped with the independently ranked top 50 accessions based on LC-MS total oil content, demonstrating that additive stacking of favorable alleles yields measurable gains in oil content and identifies promising parental accessions for high-oil sorghum breeding. The candidate genes associated with these loci are distributed across key steps of carbon flux, fatty acid synthesis, TAG assembly, and lipid storage within the FA→TAG metabolic pathway (Fig. 5C). Together, these results demonstrate that additive stacking of favorable alleles across lipid biosynthetic loci can substantially increase grain oil content, providing a practical genomic framework for developing high-oil sorghum through marker-assisted or genomic selection.

## DISCUSSION

### Targeted Breeding for Sorghum Grain Lipids

This study establishes a comprehensive genetic framework for targeted improvement of sorghum grain lipid content and composition. By integrating high-resolution UHPLC-MS lipid profiling with genome-wide association analyses, we identified key genomic regions, haplotypes, and SNP markers that collectively provide an actionable roadmap for breeding programs. By resolving oil variation at the level of chemically defined lipid species rather than bulk fat content, this work extends grain-quality GWAS in cereals into the ‘molecular phenotype’ space, bridging quantitative genetics with lipid biochemistry in a way that has only recently been achieved for other complex traits. For e.g., In maize and rice, metabolome-scale GWAS have been used to dissect natural variation in grain metabolites and link them to genetic loci, illuminating the complex architecture of grain biochemical traits (CHEN *et al*. 2014; WEN *et al*. 2014). However, sorghum lacks a similar resource for grain lipid genetics, and our results help fill this gap while aligning sorghum with advances already made in other cereals. The integration of high-density SNP data with UHPLC-MS lipid phenotypes provides a template for future sorghum multi-omics studies that seek to connect regulatory alleles, pathway flux, and end-use quality traits within a single mapping framework.

Unlike prior GWAS that used near-infrared (NIR) spectroscopy to estimate bulk grain fat content in the SAP and identified major loci such as one on chromosome 2 near 57.7 Mb, our study dissects genetic variation at the level of individual lipid species using UHPLC-MS (RHODES *et al*. 2017). The concordance between sorghum lipid-associated loci and biosynthetic pathways implicated in maize and soybean seed oil GWAS suggests that a partially conserved genetic toolkit underlies oil accumulation across cereals and oilseeds, opening opportunities for comparative fine-mapping and cross-species candidate validation. NIR approaches capture aggregate oil phenotypes but lack resolution to link SNPs to specific triacylglycerols (TAGs), phospholipids, or sterol esters, limiting insights into acyl chain composition and metabolic pathways (RHODES *et al*. 2017). By contrast, our study resolves fine-scale allelic effects on defined lipid classes, enabling precise candidate gene prioritization and haplotype-based breeding that transcends the coarse traits available from NIR.

The five lipid-enriched genomic regions identified in this study represent core targets for sorghum breeding. In maize, a GWAS of 368 diverse inbreds uncovered 74 loci significantly associated with kernel oil concentration and fatty acid composition, explaining a substantial proportion of phenotypic variation and highlighting the polygenic basis of oil traits (LI *et al*. 2013). Complementary pathway analyses further showed that acyl-lipid biosynthesis and related pathways contribute to natural variation in maize lipid content (LI *et al*. 2019). These findings in maize demonstrate that both major and minor loci can be harnessed for breeding improvement. In contrast, oil-related GWAS in soybean have identified loci associated with grain oil content and composition, providing markers for selection in this major oilseed crop (HWANG *et al*. 2014). Whereas metabolite-scale GWAS in maize and rice have primarily targeted primary metabolites and bulk kernel composition, our findings establish grain lipid architecture in sorghum as similarly polygenic yet amenable to dissection through haplotype-aware models. Consistent with these studies, our five genomic regions in sorghum collectively account for meaningful variation in grain lipid profiles and are therefore promising targets for marker development (LI *et al*. 2019).

The 12-core haplotype-tagging SNPs within these regions provide a concise marker set for marker-assisted selection (MAS). The definition of multilocus, lipid- favorable haplotypes move beyond single- marker associations toward haplotype-based breeding; - a strategy increasingly recognized as essential for capturing local linkage and epistatic effects in complex trait improvement. These markers enable breeders to identify high-oil donor accessions in public germplasm collections and incorporate lipid-associated alleles into genomic prediction models through weighted polygenic scores. Such strategies have been effective in maize, where large-effect loci such as *ZmDGAT1-2* have been associated with oil concentration and can be selected to increase grain oil content (LI *et al*. 2013; ZHAO *et al*. 2025). Similarly, in soybeans, GWAS have guided the identification of candidate loci for grain oil and protein content, demonstrating the utility of genetic markers to facilitate breeding (HWANG *et al*. 2014). Because the 12 haplotype-tagging SNPs jointly summarize the major lipid-favorable haplotypes, they can be readily incorporated into low-density breeder genotyping platforms or imputed from existing genome-wide arrays to support routine deployment in sorghum improvement pipelines.

Importantly, our haplotype analysis provides a model of haplotype stacking-based breeding strategy. Pyramiding favorable alleles across multiple loci has proven effective in increasing grain oil concentrations in other crops (FANG *et al*. 2021). Meta-analyses in maize have emphasized the importance of combining multiple QTL and candidate genes to capture more genetic variance for oil traits, providing stable mQTL for breeding (ZHAO *et al*. 2025). The catalog of 27 sorghum accessions carrying three or more favorable haplotypes provides a practical parental panel for hybrid development and pre-breeding efforts aimed at elevating grain oil content. Strategic stacking of favorable haplotypes across the five major genomic regions is expected to generate additive gains in oil accumulation and may also enable targeted modification of lipid classes or acyl chain composition. Because the identified loci influence both total oil accumulation and specific lipid species, breeders can tailor selection schemes to meet distinct end-use goals, including nutritional quality, feed efficiency, and bioenergy applications.

Although sorghum grain oil content is generally lower than that of maize and higher than that of rice, its environmental adaptability provides strong justification for targeted oil enhancement. By translating genome-scale lipid associations into deployable markers and well-defined haplotypes, this study advances molecular breeding toward greater precision and real-world applicability. Collectively, these findings establish a practical and scalable framework for enhancing grain oil content and composition, integrating insights from other cereal crops while capitalizing on sorghum’s distinct agronomic advantages. In practical terms, these markers enable a tiered selection scheme in which early-generation lines are enriched for favorable lipid haplotypes by marker-assisted selection, while later cycles exploit genomic prediction to capture the remaining polygenic variance in grain oil traits. Future work that overlays these lipid-associated loci with transcriptomic and epigenomic maps across grain development will be critical for refining causal genes.

### Post-GWAS gene function studies

By linking hundreds of lipid-associated traits to annotated metabolic and regulatory genes, this study provides a resource for post-GWAS functional dissection of sorghum grain lipid biology. A key conceptual advance of this work is the application of multivariate GWAS to exploit covariance among lipid species. Similar metabolite-GWAS efforts in maize and rice have demonstrated that combining dense metabolomics with genomic data accelerates candidate gene discovery and pathway resolution (CHEN *et al*. 2014; WEN *et al*. 2014). In maize, metabolite-based GWAS revealed extensive network connectivity and pleiotropy across biochemical pathways by jointly analyzing correlated metabolites (WEN *et al*. 2014) illustrating how modeling trait covariance increases power to detect biologically meaningful loci. Consistent with this principle, our multivariate framework uncovered loci that were effectively invisible to single-trait scans, including candidates involved in core fatty acid biosynthesis and phospholipid trafficking pathways. These findings reinforce the concept that grain lipid metabolism is organized into co-regulated modules whose genetic effects may only emerge when trait covariance is explicitly modeled.

Despite these advances, our conclusions remain association-based and leave mechanistic questions unresolved. The causal genes within terpene and saccharide-terpene clusters that influence TAG and DAG homeostasis have yet to be experimentally validated. Moreover, it is unclear how these loci interact with canonical lipid enzymes such as DGAT1, FAD2, LACS2, and members of the KCS family-genes repeatedly implicated in oil biosynthesis and fatty acid modification in other plants (MILLAR *et al*. 1999; ZHANG *et al*. 2009; CHOUDHARY AND MISHRA 2021; WU *et al*. 2024). An especially compelling example is a lead SNP on chromosome 2 (rs2357316, Cluster 4, ∼58 Mb) located within *Sobic.002G184500*, annotated as a dual serine/threonine protein kinase-alpha amylase. This region co-localizes with a previously reported grain fat and kernel composition QTL near 57-58 Mb, where an AMY3-like alpha amylase was proposed as the causal gene (RHODES *et al*. 2017). Alpha amylases are the major starch-degrading enzymes in cereal grains and play key roles in starch turnover and carbon remobilization during grain development (WHAN *et al*. 2014). Our association suggests that alpha amylase-mediated starch turnover may represent a conserved mechanism linking carbohydrate remobilization to oil accumulation in sorghum grain. Functional comparisons of rs2357316 haplotypes for alpha amylase expression, enzyme activity, and carbon partitioning during endosperm maturation will be essential to determine whether this locus modulates the trade-off between starch and lipid storage.

Addressing these gaps will require systematic post-GWAS validation. Embryo-specific expression datasets and existing mutant resources in sorghum and maize can be combined with our association signals to prioritize high-confidence candidates for CRISPR/Cas-based knockouts, allele swaps, and transgenic complementation. For example, recent sorghum oil engineering, where the over expression of WRINKLED1, Cuphea DGAT, and oleosin transgenes boosted triacylglycerol accumulation up to 5.5% leaf dry weight in field trials (PARK *et al*. 2025).

Finally, the current SAP panel captures only part of global sorghum diversity and was evaluated under limited environmental conditions. Expanding lipidome-GWAS to broader germplasm collections-including wild relatives and specialty high-oil accessions-and conducting multi-location trials will be essential to resolve genotype x environment interactions. Integrating this lipidome atlas with expression QTL, chromatin conformation maps, and metabolite-protein interaction data will ultimately enable construction of causal gene networks connecting natural sequence variation to embryo lipid phenotypes. Together, these post-GWAS efforts will move the field from statistical associations toward a mechanistic, experimentally validated framework for engineering grain oil content and stability in sorghum and related cereals.

## MATERIALS AND METHODS

### Sample Collection

A total of 400 accessions from the Sorghum Association Panel (SAP) were obtained from the Germplasm Resources Information Network (GRIN). Plants were cultivated at the Texas Tech University Research Farm in Lubbock, Texas, USA (33°35′52.9″ N, 101°54′21.4″ W; elevation 992 m). The site is characterized by a semi-arid climate, receiving an average annual precipitation of approximately 469 mm, and features fine-loamy, mixed, superactive, thermic Aridic Paleustalf soils typical of the Amarillo series. Supplemental irrigation (∼2.56 cm week⁻¹) was applied throughout the growing season to ensure consistent plant development. To minimize outcrossing, panicles were covered by pollination bags before flowering. For lipidome profiling, a subset of 266 accessions was randomly selected based on their ability to reach physiological maturity under the local growing conditions and grain availability. Three biological replicates per accession (approximately 20 grains per replicate) were used for lipidome analysis. Grains were freeze-dried using a benchtop freeze dryer (Labconco FreeZone 2.5 L, -84°C) connected to a vacuum pump (Labconco Model 117) to prevent degradation and minimize heat generation during processing. The freeze-dried grains were then finely ground in 2 mL microcentrifuge tubes using a bead mill homogenizer.

### UHPLC-MS-Based Lipidomics

Approximately 35 mg ±10% powder from each replicate was weighed into 2 mL microcentrifuge tubes with two ceramic zirconium oxide beads added. A total of 875 μL of pre-cooled extraction solvent (methyl tert-butyl ether (MTBE): methanol (MeOH), 3:1, v/v) and 20 μL of SPLASH Lipidomic internal standard mixture were added to each sample.

Samples were homogenized using a BioSpec Mini-BeadBeater-96 at 2400 rpm for 2 × 45s. The homogenates were then incubated on an orbital shaker at 100 rpm for 45 min at room temperature (RT). To promote phase separation, 570 μL H₂O:MeOH (3:1, v/v) was added to each tube. Samples were again homogenized using the Mini-BeadBeater-96 under the same conditions (2400 rpm, 2 × 45 s) and incubated on an orbital shaker at 100 rpm for 45 min at RT.After incubation, biphasic separation was obtained by centrifugation at 20,000 × g for 5 min at 4 °C. From the resulting upper organic (lipid-containing) phase, 200 μL was transferred into a labelled 1.5 mL microcentrifuge tube. The solvent was evaporated using a vacuum concentrator (SpeedVac, no heating) for 1-2 h.

Dried lipid extracts were resuspended in 500 μL acetonitrile (ACN): 2-propanol (IPA): H₂O (65:30:5, v/v) and centrifuged at 20,000 × g for 10 min at 4 °C. 120 μL of the clarified supernatant was transferred into autosampler vials for LC-MS analysis.

Chromatographic separation was performed using a Thermo Scientific Vanquish UHPLC system equipped with a Waters Acquity UHPLC BEH C8 column (1.7 µm, 2.1 × 100 mm). The column temperature was maintained at 55 °C, the mobile phase flow rate was 0.350 mL/min, and the injection volume was 10 μL.Mobile phase A consisted of 60% ACN, 40% water, 10 mM ammonium formate, and 0.1% formic acid, and mobile phase B consisted of 88% IPA, 10% ACN, 2% water, 10 mM ammonium formate, and 0.1% formic acid.

The LC gradient started at 35% B (0-4 min), increased to 60% B (4-12 min), then to 85% B (12-21 min), followed by 100% B (21-21.1 min) and held at 100% B (21.1-24 min). The gradient was then returned to 35% B at 24.1 min and maintained until 28 min.

Mass spectrometry was performed on a Q-Exactive HF mass spectrometer (Thermo Fisher Scientific) operated with electrospray ionization (ESI) using full MS data-dependent MS² (ddMS²) acquisition in both positive and negative modes. Full MS parameters were: 120,000 resolutions, AGC target 1 × 10⁶, maximum IT 100 ms, and scan range m/z 250-1200. ddMS² parameters were: 30,000 resolution, AGC target 2 × 10⁵, maximum IT 75 ms, loop count 10, isolation window 1.0 m/z, and normalized collision energy (NCE) of 25 and 30.

### Lipid identification and quantification

Lipid separation and detection were conducted on a Waters 2777c UHPLC system coupled to a Q Exactive HF high-resolution mass spectrometer (Thermo Fisher Scientific, USA), operating in both positive and negative electrospray ionization (ESI) modes. Raw MS data (.raw) were converted to .mzXML format using MSConvert (ADUSUMILLI AND MALLICK 2017). Feature detection, alignment, and quantification were performed using XCMS (SMITH *et al*. 2006) and CAMERA (KUHL *et al*. 2012), retaining features present in at least three samples to preserve low-abundance lipids. Missing values were imputed using median or interpolation-based methods. Mass features were filtered by retention time (60-630 s) and *m/z* precision (< 0.5 Da), excluding isotopic peaks while retaining MS adducts to preserve true lipid features. Feature intensities were normalized to tissue fresh weight and total ion current to correct for technical variability. Log-transformation and Pareto scaling were applied before statistical analysis (VAN DEN BERG *et al*. 2006). Metabolic trait (*m*-trait) data were calculated as the average of three biological replicates for each lipid (NIGAM *et al*. 2025). Lipid species were identified by matching accurate mass and retention time to databases including LIPID MAPS (O’DONNELL *et al*. 2019), HMDB (WISHART *et al*. 2022), and KEGG (KANEHISA 2002), allowing classification of triacylglycerols (TAGs), phosphatidylcholines (PCs), phosphatidylethanolamines (PEs), and other lipid classes.

Raw lipidomic data files were processed using LipidSearch software (version 5.1.6, Thermo Fisher Scientific, Waltham, MA, USA), a leading commercial lipidomics platform for lipid identification and relative quantification (YAMADA *et al*. 2013; CHONG *et al*. 2018). LipidSearch was run with an integrated database search using a mass tolerance of 5 ppm for precursor ions and 8 ppm for product ions, and an absolute intensity threshold of 30,000 (FERNÁNDEZ-LÓPEZ *et al*. 2019). Lipid identifications were restricted to species with a match score >5.0 that were detected in at least two technical replicates. Peak area integration was used for quantification, and values were normalized to internal standards prior to downstream analysis. The resulting lipid abundance matrix was exported for statistical analyses in R (version 4.5.3), and data visualization was performed using the ggplot2 package (v4.0.2) (WICKHAM 2016).

### Hierarchical Clustering of Lipid Species

The 528 lipid species were hierarchically clustered to identify groups of correlated lipids for subsequent multivariate GWAS analyses (CARLSON *et al*. 2019; WANG *et al*. 2022; POTAPOVA *et al*. 2023). Absolute pairwise Pearson correlations were calculated using inverse-normal transformed lipid levels. Clustering was conducted separately for glycerolipids (from TAGs and DAGs) and for the remaining lipid species (from glycerophospholipids). To reduce instability in multivariate association analyses caused by highly correlated traits, we iteratively removed one lipid from each pair with correlation >0.8 until no pair exceeded this threshold. Hierarchical clustering was then applied to the filtered lipid set using average linkage based on Euclidean distances derived from pairwise correlations. Clusters were visually identified from the resulting dendrogram. Within each cluster, variance inflation factors (VIFs) were calculated for every lipid species using the R package car (GROß 2003; OTTENSMANN *et al*. 2023). Cluster members were treated as independent variables in a regression model with a randomly generated outcome, as the outcome values do not affect VIF calculation (KUTNER 1984). Iteratively, the lipid species with the highest VIF was removed until all cluster members had a VIF < 5, thereby eliminating traits highly correlated with linear combinations of other cluster members.

### Lipidome-based genome-wide association study (lGWAS)

GWAS were conducted to identify genetic loci controlling variation in sorghum grain lipid composition across 266 accessions of the SAP. Published SAP SNP data aligned to the *Sorghum bicolor* BTx623 reference genome (v3.1) were filtered to retain variants with minor allele frequency (MAF ≥ 0.05) and the missing rate ≤ 10%. After quality control, 38,873,896 high-quality SNPs were retained for analysis (BOATWRIGHT *et al*. 2022). Missing genotypes were imputed using Beagle (v5.0) with default parameters. Genome-wide linkage disequilibrium (LD) decay was estimated using PopLDdecay (ZHANG *et al*. 2019), revealing an average LD decay distance of ∼50 kb at r² = 0.2. This decay distance was used to define independent association loci.

Lipid abundances were log₁₀-transformed and inverse-normal transformed prior to association testing to improve normality and reduce heteroscedasticity. Only lipid species detected in ≥75% of accessions were retained, resulting in 528 lipid traits for analysis. A univariate GWAS was conducted for each lipid species using a mixed linear model (MLM) implemented in the R package rMVP (v1.0.1) (YIN *et al*. 2021). Population structure was controlled by including the first three principal components derived from genome-wide SNPs. The kinship matrix was calculated using the VanRaden method and incorporated as a random effect (VANRADEN 2008).

We conducted GWAS using the multi-locus model FarmCPU (Fixed and Random Model Circulating Probability Unification) implemented in rMVP (LIU *et al*. 2016).

FarmCPU was run with the first three PCs as covariates, maxLoop = 10, method.bin = “FaST-LMM,” and variance components estimated using the “GEMMA” option. To assess robustness, FarmCPU was repeated 100 times using 90% random subsampling, and Resample Model Inclusion Probability (RMIP) was calculated for each SNP. Only SNPs with RMIP > 0.05 were considered stable associations (VALDAR *et al*. 2009).

To leverage correlations among lipid species and increase power to detect pleiotropic loci, multivariate GWAS was performed by jointly modeling six biologically defined lipid clusters: triacylglycerols (TAGs), phosphatidylcholines/phosphatidylethanolamines (PCs/PEs), sphingolipids, galactolipids, phosphatidylinositols, and minor lipid classes. The multivariate framework estimated posterior inclusion probabilities (PIP) and local false sign rates (LFSR) to quantify the probability and direction of SNP effects across correlated lipid traits.

Genome-wide significance thresholds were determined using Bonferroni correction based on the effective number of independent markers estimated using the Genetic Type I Error Calculator (GEC v0.2) (LI *et al*. 2012). A conservative threshold of *P* <10^−8^ was applied to control the family-wise error rate. Quantile-quantile (QQ) plots were inspected to verify appropriate control of population structure, with genomic inflation factors ranging from 0.98 to 1.15 (YANG *et al*. 2011). Significant SNPs were grouped into independent loci by merging variants within ±50 kb windows, consistent with the observed LD decay. Lead SNPs were defined as the most significant variant within each locus after accounting for LD structure (r² ≥ 0.1). Candidate genes within each locus were annotated using sorghum BTx623 reference genome (v5.1, Phytozome) (GOODSTEIN *et al*. 2012).

### Identification of lipidome-linked variants in metabolic gene clusters

Metabolic gene clusters (MGCs) in sorghum were identified using the plantiSMASH web platform (version 2.0-beta4) following standard analytical workflows (KAUTSAR *et al*. 2017). A CD-HIT clustering threshold of 0.5 was applied to group proteins sharing ≥ 50% sequence identity (HUANG *et al*. 2010), and clusters containing a minimum of two unique biosynthetic domains were retained (GRAßMANN 2005). Built-in profile Hidden Markov Models (pHMMs) were used to enhance the recognition of plant-specific biosynthetic motifs (FRIEDRICH *et al*. 2006). Predicted clusters were manually curated to confirm domain composition. To assess the genomic association between lipidome-linked variants and MGCs, SNPs identified through lGWAS were mapped to the MGC coordinates. Variants within ±50 kb and ±500 kb flanking regions of each cluster were quantified, and enrichment ratios (observed/expected) were computed based on random permutation tests (10,000 iterations) to evaluate statistical significance (*P* < 0.05). SNPs were further categorized by lipid class-including triacylglycerols (TAG), phosphatidylglycerols (PG), phosphatidylcholines (PC), and phosphatidylethanolamines (PE)-and visualized using bar plots and dot plots to illustrate class-specific enrichment patterns and allele distributions across selected clusters.

### Polygenic Lipid Score (PLS) Analysis

To quantify the cumulative effect of oil-associated loci, a Polygenic Lipid Score (PLS) was computed for each sorghum line in the population. The PLS integrates additive effects (POWELL *et al*. 2013) estimated from metabolic GWAS (mGWAS) and multivariate analyses, capturing the aggregate contribution of favorable alleles across the genome. For each significant SNP, the estimated allelic substitution effect (*β_j_*) was multiplied by the genotype dosage (*g_ij_*; coded as 0, 1, or 2 according to the number of favorable alleles). The total PLS for each line was calculated as:

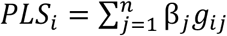

where *PLS_i_* is the polygenic lipid score of line *i* and *n* is the number of independent oil-associated loci included in the model. To minimize redundancy from linkage disequilibrium (LD), only lead or conditionally independent SNPs were retained. All PLS values were standardized to have a mean of zero and a standard deviation of one before further analysis. The relationship between the PLS and measured oil content was modeled using a simple linear regression framework (COLLISTER *et al*. 2022; ABU-EL-HAIJA *et al*. 2023):

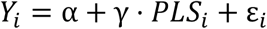

where *Y_i_* denotes the measured oil content of line *i*, *α*is the intercept, *γ*represents the regression coefficient describing the effect of the PLS, and *ε_i_* captures residual variation. Model performance was assessed by the coefficient of determination:

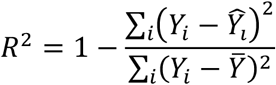

To visualize the cumulative impact of beneficial alleles, accessions were divided into five PLS quantiles (Q1-Q5) according to their standardized scores, and mean oil content was compared across quantiles as:

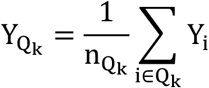

where *Ȳ_Qk_* is the mean oil content of accessions in quantile *Qk* (k = 1…5). Differences among quantiles were tested by one-way ANOVA followed by Tukey’s HSD post hoc comparisons, with *p* < 0.05 considered statistically significant.

## Author contributions

YJ conceptualized and designed the project. SM, and FC conducted lipidome profiling. DN performed data analyses. DN and YJ drafted the manuscript. All authors reviewed, edited, and approved the final version of the manuscript.

## Acknowledgments

This research was supported by the Non-Assistance Cooperative Agreement 58-3020-2-024 between Texas Tech University and USDA-ARS (3020-43440-002-00D). The authors are grateful to the publicly available USDA Germplasm Resources Information Network (GRIN) database for providing accessions of the SAP used in this study.

## Supplementary figures

**Supplemental Figure 1. Lipid composition and correlation structure of the sorghum grain lipidome.** (A) Distribution of identified lipid species across major lipid classes. Bars represent the number of detected lipid species within each class, including phosphatidylcholine (PC), phosphatidylethanolamine (PE), phosphatidylglycerol (PG), phosphatidylinositol (PI), triacylglycerols (TAG), diacylglycerols (DAG), digalactosyldiacylglycerols (DGDG), and monogalactosyldiacylglycerols (MGDG). (B) Pairwise Pearson correlation matrix of lipid abundances across 266 sorghum accessions. Lipids were hierarchically clustered based on correlation coefficients. The color scale represents Pearson correlation values ranging from negative to positive correlations. Colored side bars denote lipid clusters identified for multivariate analysis.

**Supplemental Figure 2. Principal component analysis of sorghum grain lipid profiles.**(A) Proportion of variance explained by the first ten principal components derived from lipid abundance data across 266 sorghum accessions. (B) Principal component analysis (PCA) biplot showing the distribution of sorghum accessions based on lipid composition. Each point represents an accession, and the axes correspond to the first two principal components (PC1 and PC2). Arrows represent lipid species loadings, with arrow length indicating relative contribution and direction corresponding to loading orientation in the ordination space.

**Supplemental Figure 3. Hierarchical clustering and principal component analysis of sorghum grain lipid species.** (A) Heatmap of normalized lipid abundances across 266 sorghum accessions. Rows represent individual lipid species and columns represent sorghum accessions. Both lipids and accessions were hierarchically clustered based on similarity in lipid abundance profiles. Colored side bars denote lipid classes. (B) Principal component analysis (PCA) of lipid species based on scaled abundances. Each point represents a lipid species colored by lipid class, and point size corresponds to mean log₁₀ abundance. The percentages on the axes indicate the proportion of variance explained by PC1 and PC2. (C) Abundance-weighted principal component analysis (PCA) of lipid species based on weighted abundance values.

**Supplemental Figure 4. Genome-wide distribution of lipid-associated loci and comparison of univariate and multivariate GWAS results.** (A) Genome-wide distribution of significant SNP associations for major lipid classes across sorghum chromosomes. Points represent significant associations for triacylglycerols (TAG), phosphatidylethanolamines (PE), and phosphatidylcholines (PC), plotted according to chromosomal position. (B) Comparison of association significance between univariate and multivariate GWAS analyses. Each point represents an association region colored by lipid class. The x-axis shows −log₁₀(p) values from univariate GWAS and the y-axis shows −log₁₀(p) values from multivariate GWAS. The diagonal line indicates equal −log₁₀(p) values between the two approaches.

**Supplemental Figure 5. Enrichment of lipid-associated SNPs in specialized Metabolic Gene Clusters (MGCs).** (A) Distribution of GWAS-significant SNPs located within ±50, ±100, and ±500 kb of specialized metabolic gene clusters. Bars indicate the number of lipid-associated SNPs across different cluster types and cluster IDs. (B) Enrichment analysis of lipid-associated SNPs within metabolic gene clusters. Bars represent the ratio of observed to expected SNP counts for each cluster at different window sizes. Asterisks denote clusters meeting the statistical significance threshold. (C) Genomic locations of lipid-associated SNPs within selected metabolic gene clusters. Colored points represent SNPs associated with different lipid classes (TAG, PC, and PE), and gene models are shown along genomic coordinates.

**Supplemental Figure 6. Functional enrichment and metabolic pathway mapping of lipid-associated candidate genes. (A)** Functional enrichment analysis of candidate genes located near lipid-associated loci. Enriched biological pathways are shown on the y-axis, and the enrichment metric is shown on the x-axis. Circle size represents the number of genes associated with each pathway, and color indicates the false discovery rate (FDR). **(B)** Mapping of candidate genes onto the glycerolipid metabolic pathway. Highlighted enzyme steps correspond to reactions associated with candidate genes identified near lipid-associated loci.

**Supplementary Figure 7. Overview of the analytical framework for identifying and stacking favorable lipid-associated alleles.** Workflow illustrating lipidomic profiling of 266 sorghum accessions using UHPLC-MS to quantify lipid traits and define lipid clusters. Univariate and multivariate GWAS analyses were conducted to identify lipid-associated loci. Significant loci were integrated to construct a Polygenic Lipid Score (PLS), calculated as the weighted sum of favorable allele dosages based on GWAS effect sizes. The PLS was incorporated into a regression model to assess its relationship with total oil content.

## Supplementary files

**Supplementary Table S1. Lipidomic species identifiers and normalized peak areas across sorghum grain accessions.** Lipid species detected in sorghum grain using LC-MS analysis. Each row represents a lipid feature with annotation information including mass-to-charge ratio (m/z), retention time, compound identification, lipid classification, and database identifiers. Additional columns provide peak detection quality metrics and normalized peak areas for each sorghum germplasm accession (PI numbers), representing the relative abundance of each lipid species across the analyzed accessions.

**Supplementary Table S2. Lipid species from six hierarchical clusters in the sorghum grain lipidome.** Lipid species identified in sorghum grain and grouped into six clusters based on hierarchical clustering of lipid abundance profiles across sorghum accessions. The table includes the lipid name, lipid class, and the corresponding cluster assignment.

**Supplementary Table S3. Lipidome PCA variance and cluster statistics. PC loadings and ANOVA for high- and low-oil sorghum groups**. Principal component analysis (PCA) scores for sorghum grain accessions based on lipidomic profiles. The table includes PC1-PC4 scores, cluster assignments derived from lipidome-based grouping (e.g., high-oil and low-oil clusters), and identification of accessions belonging to the top 20 high-oil group. These results summarize lipidome variation among sorghum accessions and highlight clusters associated with elevated grain oil content.

**Supplementary Table S4. Genomic control inflation factor (λ) for individual lipidome traits used in GWAS analysis.** Lipid species detected in the sorghum grain lipidome along with their annotation information, including mass-to-charge ratio (m/z), retention time, compound identification, chemical formula, lipid classification, and database identifiers. The table also reports the genomic control inflation factor (λ) calculated for each lipid trait to assess the level of test statistic inflation in genome-wide association analysis.

**Supplementary Table S5. Lead SNP clusters, nearby genes within ±50 kb genomic windows, and association statistics for each lipid species across six LD-defined clusters (r² ≥ 0.1).** Lead SNPs identified from GWAS of sorghum grain lipid traits, grouped into LD-defined clusters. The table includes SNP genomic positions, nearby genes within ±50 kb windows, gene annotations, allele information, minor allele frequency (MAF), and association statistics estimated using the FarmCPU model.

**Supplementary Table S6. Predicted biosynthetic gene clusters associated with lipid pathways.** Genomic regions predicted to contain lipid-related biosynthetic gene clusters, including genomic coordinates, core biosynthetic domain annotations, similarity to known clusters, and counts of nearby SNPs within 50 kb, 100 kb, and 500 kb genomic windows.

**Supplementary Table S7. Functional enrichment analysis of proteins in the interaction network.** Enriched metabolic pathways identified from proteins in the network, including pathway names, gene counts, enrichment strength, and corresponding false discovery rate (FDR) values.

**Supplementary Table S8. Germplasm lines ranked by polygenic lipid score (PLS) and predicted lipid content.** Sorghum germplasm accessions ranked based on their polygenic lipid score (PLS) and predicted lipid content, including lipid level classification, counts of beneficial alleles, and log₁₀-transformed values of PLS and predicted lipid content.

